# Unexpected mixed-mode transmission and moderate genetic regulation of *Symbiodinium* communities in a brooding coral

**DOI:** 10.1101/173591

**Authors:** Kate M. Quigley, Patricia A. Warner, Line K. Bay, Bette L. Willis

## Abstract

Determining the extent to which *Symbiodinium* communities in corals are inherited versus environmentally-acquired is fundamental to understanding coral resilience and to predicting coral responses to stressors like warming oceans that disrupt this critical endosymbiosis. We examined the fidelity with which *Symbiodinium* communities in the brooding coral *Seriatopora hystrix* are vertically transmitted and the extent to which communities are genetically regulated, by genotyping 60 larvae and their parents (9 maternal and 45 paternal colonies) using high throughput sequencing of the ITS-2 locus. Unexpectedly, *Symbiodinium* communities associated with brooded larvae were distinct from those within parent colonies, including the presence of types not detected in adults. Bayesian heritability (h^2^) analysis revealed that 33% of variability in larval *Symbiodinium* communities was genetically controlled. Results highlight flexibility in the establishment of larval communities and overturn the paradigm that symbiont transmission is exclusively vertical in brooding corals. Instead, we show that *Symbiodinium* transmission in *S. hystrix* involves a mixed-mode strategy, similar to many terrestrial invertebrate symbioses. Also, variation in the abundances of common *Symbiodinium* types among adult communities suggests that microhabitat differences influence the structure of *in hospite Symbiodinium* communities. Partial genetic regulation coupled with flexibility in the environmentally-acquired component of larval *Symbiodinium* communities implies that corals with vertical transmission, like *S. hystrix,* may be more resilient to environmental change than previously thought.

## Introduction

Symbiosis is fundamental to life on Earth, underpinning the existence of numerous prokaryotic and eukaryotic species and shaping the physiology and health of many organisms [1–3]. Microbial symbionts also enable hosts to expand their niche breadth to survive in environments otherwise unsuited to their physiology [4]. For example, symbiosis with photosynthetic dinoflagellates of the genus *Symbiodinium* has allowed corals to thrive in oligotrophic tropical seas through the utilization of symbiont photosynthates. Similar nutritional facilitation has been described for sap-sucking insects that rely on microbial partners to supplement their diets [5] and legumes that rely on rhizobia to fix nitrogen [6]. Unlike these well-characterized systems, coral endosymbioses are poorly described at the *Symbiodinium* type level during early ontogeny.

Nutritional symbioses can drive diversification of host and symbiont lineages [7–9], with eukaryotic symbionts like *Symbiodinium* having gone through multiple cycles of diversification and expansion [10]. Such sources of genetic variation provide new material upon which selection may operate [11,12], facilitating coevolution between hosts and symbionts or among symbionts [3,11,13]. Understanding the fidelity (exactness of transfer of symbionts from parent to offspring) of *Symbiodinium* community inheritance is key to determining the degree to which endosymbiotic *Symbiodinium* communities have coevolved with their coral hosts and is central to coral nutrition and health. Despite this, little is known about genetic regulation underpinning this symbiosis.

Symbionts may be acquired from the environment (horizontal transmission) or passed maternally into eggs or larvae (vertical transmission), with the latter thought to be the most prevalent mode of transmission in brooding scleractinian corals [14]. Maternally-derived symbionts may involve the transmission of one or multiple symbionts (superinfections) and, at least in well-studied insect vertical symbioses, may strongly impact host reproduction, behaviour and co-evolution [12,15]. Transmission of insect symbionts may be exclusively vertical or may occur initially as vertical transfer followed later by horizontal transmission [9,15–19]. Although similar mixed-mode transmission has been hypothesized for corals [20], the absence of experimental data means that it is not yet clear if transmission is exclusively vertical in brooding corals or if mixed-mode transmission also occurs in this group. Given recent evidence of differences in the diversity of symbiont communities transmitted from parents to offspring in two broadcast spawning corals [21,22], *Symbiodinium* transmission dynamics may be as complex as those observed in the Arthropoda.

In general, symbiont-host specificity is theorized to be much greater when symbionts are transmitted vertically compared to horizontally [23,24]. In corals, hosts may form strict associations with only one *Symbiodinium* type (and *vice versa*) or associate with multiple partners, but in general, superinfections of multiple *Symbiodinium* types and subtypes of varying abundances are common [20,25–29]. Although maternal transfer of *Symbiodinium* and bacteria is less well-characterized in corals than in terrestrial invertebrates [20–22,30,31], the presence of superinfections raises the possibility that *Symbiodinium* dynamics are similar to the mixed-mode transmission dynamics characteristic of superinfections described in terrestrial invertebrates like aphids and sharpshooter cicada [9,11]. However, unlike studies of insect symbiont specificity, no studies have used high throughput sequencing to examine maternally-transmitted *Symbiodinium* communities in brooding corals or the diversity of low abundance *Symbiodinium* types in detail. Similarly, the genetic component of parental contributions to the maturation of *coral*-*Symbiodinium* symbioses remains unquantified.

It is clear that different *Symbiodinium* types vary in their impact on holobiont physiology because of variation in their stress tolerance and ability to produce and transfer photosynthates to the coral host under differing light, temperature and nutrient regimes [28,32–36]. Moreover, environmental stressors may bring about shifts in the dominance of different *Symbiodinium* types, in some cases benefiting the host under the altered conditions [37,38]. The extent of a coral’s flexibility to acquire resilient types or shuffle symbionts may be genetically regulated, for example by heritable host immune responses, similar to those that shape symbiont diversity in *Drosophila* [39]. Complete inheritance results in complete fidelity of symbiont transmission, and, hence, little scope for flexibility in coral-*Symbiodinium* symbioses. However, the extent of such potential regulation of symbiont transmission and its underlying basis are unknown for corals.

Increasingly, studies are revealing that the genetic architecture behind traits and pathologies can be complex [40]. For example, both the diversity and abundance of microbial symbionts in the human gut are complex traits under partial genetic control [41–46]. Narrow-sense heritability (h^2^) is the parameter typically used to describe the degree to which variability in a trait is explained by genetic factors. Assuming that the *Symbiodinium* community associated with a coral can be represented as a complex trait, then an h^2^ value of 1 would imply that variability of the community is mostly due to host genetics. Conversely, an h^2^ value estimated at 0 would imply no genetic basis for variability in the community, thus the community would not be under selection and could not evolve [no evolvability; 47]. Although an h^2^ estimate close to 1 does not necessarily guarantee absolute genetic determination as a result of gene segregation [48], a large heritability estimate of the *Symbiodinium* community would imply that changes in host genotypes are required for shifts in symbiont communities. Conversely, changes in the environmental availability of *Symbiodinium* or in environmental conditions would have limited influence on *in hospite* communities. Understanding the relative contributions that host genetics versus environmental conditions make to the composition of *Symbiodinium* communities through estimations of h^2^ will improve the accuracy with which the potential, direction and speed of changes in *Symbiodinium* communities can be predicted.

To examine *Symbiodinium* community transfer between adults and their offspring in a brooding coral and quantify the narrow-sense heritability (h^2^) of this trait, we quantified the *in hospite Symbiodinium* communities of individual planula larvae and their parents across a spectrum of relatedness using high throughput sequencing. Relatedness was based on a population genetic parentage analysis that assigned the likely paternal identity of each larva. In light of results on heritability and fidelity of symbiont transfer, we discuss the potential of larvae from brooding corals like *S. hystrix* to acclimate to novel environments.

## Results

### Symbiodinium communities differ between parents and brooded larvae

*Symbiodinium* communities differed between adults and their larvae in the brooding coral *Seriatopora hystrix* (ShA) (Figure 1A, B). Overall, the composition of *Symbiodinium* communities was similar among adult corals, but more variable among larvae. On average, adults contained 29.9±0.6 (SE) OTUs and larvae had 22±0.4 OTUs. However, the number of unique OTUs recovered was more than five times greater from larvae than from adults (93 vs. 17 OTUs, respectively; Figure 1C). Of the 17 unique adult OTUs, ten belonged to clade C (C1, C15 and other variants), three from clade A (A1 and variants), three from D (including D1 and D1a), and one was a putative C type (Figure 1B, sequences most highly similar to mixed *S. hystrix/Symbiodinium* libraries– see Supporting Information). Unique to larvae were 17 OTUs from clade C (likely C1 variants), four from clade E, one from each of A3, B1, and G6, and 69 that were of putative *Symbiodinium* identity (Figure 1B). Of the 93 larval-specific OTUs, only the abundance of C1_OTU136 (type followed by OTU designation) and two putative clade D OTUs (OTU148 and OTU149) were significantly different from zero with the Bejamini-Hochberg correction (Figure 1B). Although raw read counts were low, C1_OTU136 was present in larvae from every dam but dam 3.

**Figure 1.**
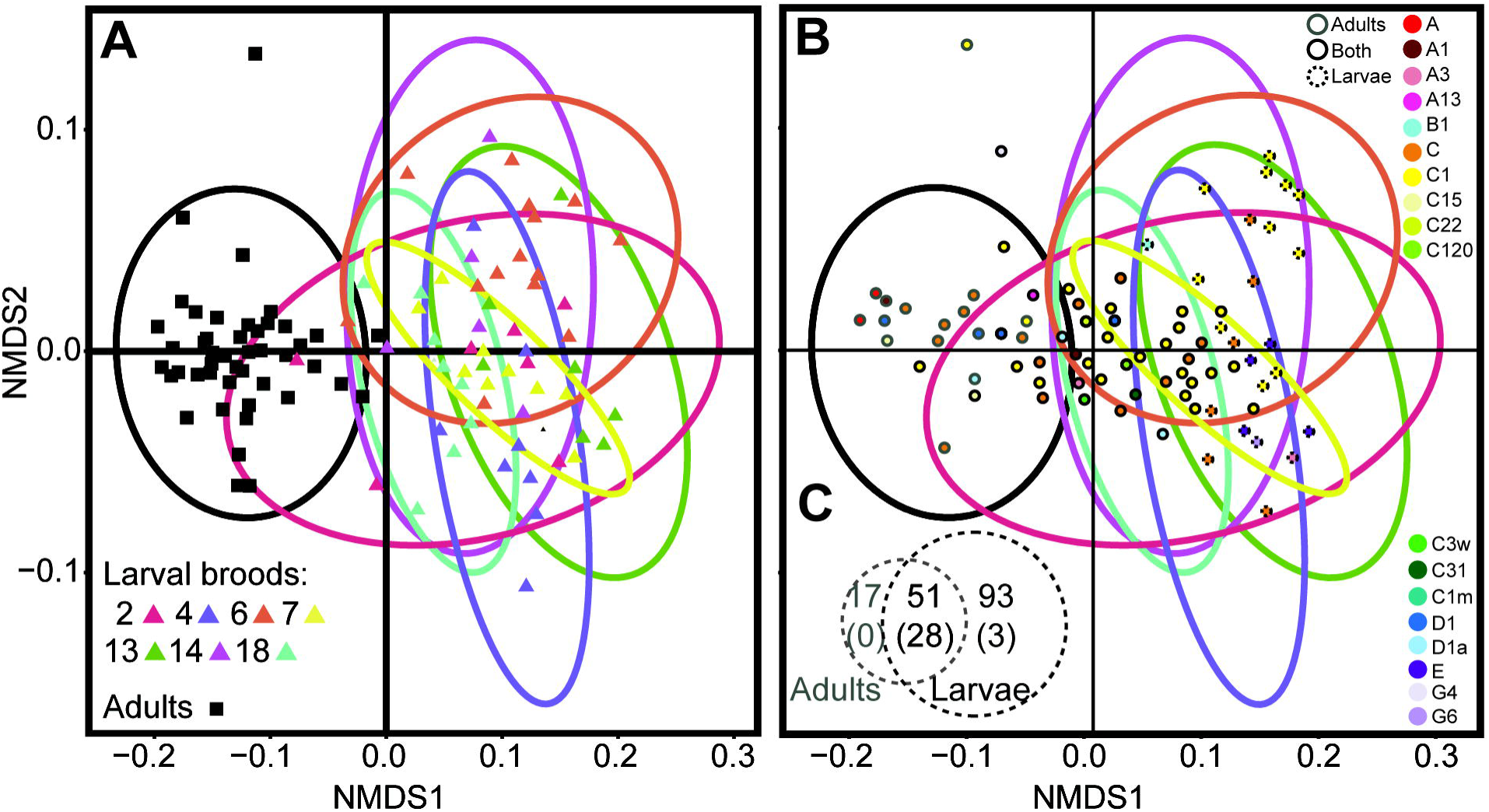
Nonmetric multidimensional scaling (NMDS) plots, based on a Bray-Curtis distance matrix of variance-normalized OTU abundances and sequence similarity between OTUs (pairwise percent identities), illustrating differences between *Symbiodinium* communities associated with adult colonies and larvae of the brooding coral *Seriatopora hystrix* (ShA). Ellipses encircling symbols of the corresponding colour represent 95% probability regions for adults (black) and larval broods (coloured), where each brood represents all larvae sharing the same dam (colour-coded). A) Each point represents the *Symbiodinium* community associated with a unique coral adult or larval sample. B) Each point represents an OTU coloured by type level (see Supporting Information table S5 for full names). Outlining around each point represents the origin of the OTU, i.e., those found uniquely in adult (grey outline) or larval (broken grey outline) samples, or retrieved from both (black outline). Samples presented in (A) and OTUs presented in (B) share the same ordination space but were separated for clarity. C) Venn diagram, illustrating the number of *Symbiodinium* OTU’s that were unique to larvae (dark grey text) versus adults (light grey text). The number of OTUs that were significant after p-adjustments are in parentheses. Ellipses corresponding to dams 3 and 10 are not represented, as only one larva per dam was collected and sequenced.

Fifty-one OTUs were shared by adult colonies and planula larvae (43 of known *Symbiodinium* taxonomy, 8 putative *Symbiodinium,* Figure 2B), and the abundances of 28 of these OTUs differed significantly between the two groups at the adjusted p-level (Table S4, Supporting Information). Of these 28 OTUs, 23 were from clade C (including C1, C3w, C120 amongst others), three from clade D (D1, D1a), and two from clade A (A1, A3). Adult *Symbiodinium* communities were characterised by up to 1.3 times more D-types (D1_OTU3, D1_OTU597, D1a_OTU6), and A-types (A1_OTU10 and A3_OTU8) compared to larvae (Bejamini-Hochberg corrections, Table S4). Nine of the 23 C-types had up to 1.7 times significantly higher abundances in adults (including multiple C1 types, C120/C120a_OTU1, C1m_OTU5/105, C1v6/C22_OTU228, C15_OTU46, C31_OTU733, and C3W_OTU165) and the remaining C1 types had between 1.5 – 2.4 times significantly lower abundances in adults (Table S4).

**Figure 2.**
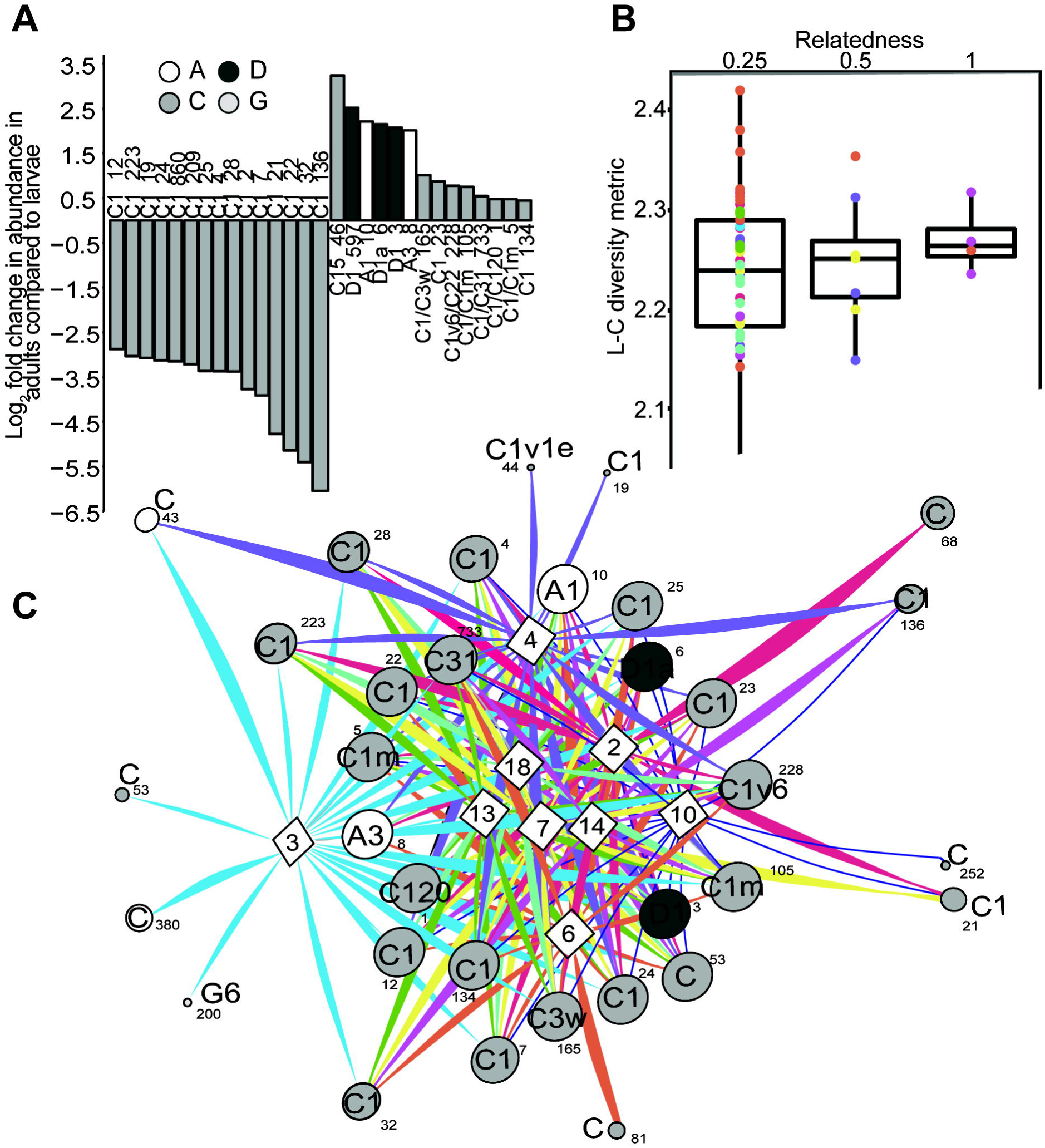
A) Log_2_ fold change in abundances of *Symbiodinium* OTU’s that differed significantly between communities associated with adults versus larvae of *Seriatopora hystrix* (ShA). Grey-scale in the bar plot identify *Symbiodinium* clades. A positive change indicates the OTU is more abundant in adults. B) Boxplots showing medians, quartiles and minimum/maximum values of *Symbiodinium* community diversity (Leinster and Cobbold metric) in relation to individual larval relatedness. On the x-axis, 0.25 denotes half sibs, 0.5 full sibs, and 1.0 denotes larvae produced from selfing. Each larva is coloured by its respective dam. C) Network analysis of planula larvae showing OTUs present in 50% or more of larvae per brood. White diamonds correspond to maternal broods, where each brood sharing the same dam is colour-coded. Line thickness denotes relative abundance of the Symbiodinium type per brood.

### Larval Symbiodinium communities vary among broods

Planula larvae that shared the same maternal parent generally clustered when *Symbiodinium* OTU richness, abundance, and DNA distance between OTUs were incorporated into analyses (Figure 1A, B). Thirty-one OTUs (including multiple C1 variants, D1, D1a, A1, and A3) were found in greater than 50% of larvae per brood and were generally present across all broods (Figure 2C). However, differences in the abundance of OTUs amongst larval broods were detected for symbiont types A1, A3, C1, D1, and D1a, amongst others. Briefly, larvae from dam 2 displayed higher abundances of A1 and A3. Larvae from dams 3, 7 and 10 had significantly less of C1_OTU2, whilst broods from dams 4, 6, 13, 14, and 18 had significantly different abundances of many C-types, including C120/C120a, C1, C1v1e, C1m, and C31. The abundances of D1_OTU3 and D1a_OTU6 also varied significantly among larval broods, particularly among those from dams 2, 4, and 18 vs. dam 13 (for a full description see Supporting Information and Table S4).

### Heritability

Leinster and Cobbold estimates of *Symbiodinium* community diversity varied across the 60 larvae. Notably, variance around median estimates decreased as relatedness between individual larvae increased (Figure 2B). The posterior mean heritability of the *Symbiodinium* community in *S. hystrix* (ShA) larvae was 0.43 ± 0.21 SD, with a posterior mode of 0.33 (95% Bayesian credibility interval (BCI) 0.1 - 0.8; Figure S2, Supporting Information). Adding maternal identity as a random effect did not improve the model (Deviance Information Criterion < 2 units) but decreased the posterior mean and mode of heritability slightly (mean = 0.37 ± 0.21 SD and mode = 0.19; BCI: 0.1 - 0.8).

### Patterns in adult Symbiodinium communities with colony size and spatial distribution

Of the 68 *Symbiodinium* OTUs found in adults, the abundance of only four C-type OTUs differed significantly among the five coral size classes. Two C-types (OTU228 and OTU105) had two times greater abundances in the 26 – 32 cm class compared to corals in each of the other four size classes (all p’s>0.05; Table S4). Similarly, C31_OTU733 was found at significantly lower abundance in corals from the 8 – 14 cm class compared to the single colony in the largest size class (p>0.05; Table S4). Colonies from the 8 – 14 cm class also had 1.7 times lower abundances of C1_OTU4 compared to corals in the 14 – 20 cm class (p>0.05; Table S4).

The distributions of three of the ten most abundant OTUs in adult corals varied significantly across the sampling area (*p* > 0.05; Figure 3), although not in a consistent manner with distance either along or down the sampling area. For example, although abundances of *Symbiodinium* C120/C120a were greatest in the lower left of the sampling area (Gradient Boosted Model (GBM): *p* = 0.019), consistent with a gradual increase in distance down the reef slope, this pattern was not consistent along the reef slope (GBM: *p* = 0.00841). The abundance of D1 was significantly higher in the top-right and lower left side of the sampling area than in other aspects (x and y interaction, GBM: *p* = 0.0393). D1a was least abundant in the top left and inner portion of the sampling area (GBM: *p* = 0.0405). Finally, although the variance normalized abundances of all three OTUs were significantly positively correlated overall (Spearman’s rank correlation rho: 0.42 – 0.77, all p < 0.004), extremely low abundances of C120/C120a at x-coordinates >15 contrast markedly with high abundances of the two D-types in the same region (Figure 3).

**Figure 3.**
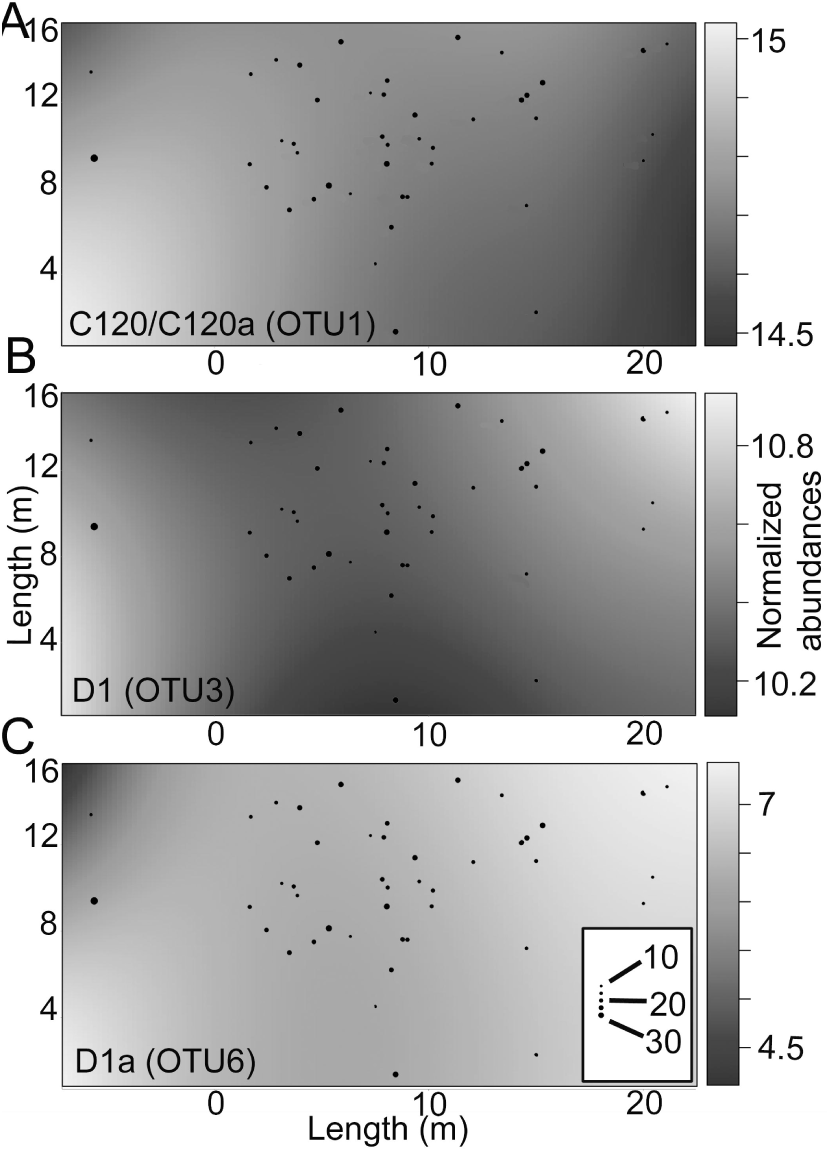
Spatial patterns in the normalised abundance of three *Symbiodinium* OTU’s associated with adult colonies of *Seriatopora hystrix* (ShA) that differed significantly in their abundances across a portion of the 16 m × 40 m sampling area at Lizard Island: A) C120/C120a, B) D1, and C) D1a. Grey scale represents changes in the normalized abundance of each OTU across sampling site coordinates. Sizes of the black circles represent size classes of coral colonies in cm (drawn to scale).

## Discussion

### Mixed mode transmission structures larval Symbiodinium communities in a brooding coral

The availability of a full larval pedigree for *Seriatopora hystrix* (ShA) [49] provided a unique opportunity to evaluate the relative contributions of heritability (i.e., the degree to which variability in a trait is explained by genetic factors) versus maternal environmental effects (the effect of larvae sharing a common maternal environment) to the composition of larval *Symbiodinium* communities in a brooding coral. Here we show that *Symbiodinium* communities associated with larvae of *S. hystrix* (ShA) differ from those associated with their parents, providing experimental evidence that at least a portion of the *Symbiodinium* community is horizontally transmitted in a brooding coral. Such paradigm-changing knowledge on symbiont transmission is important as it realigns the cnidarian literature with well-characterized models of invertebrate symbioses. Overall, *Symbiodinium* communities were found to be moderately heritable, with only 33% of variability in larval symbiont communities under genetic regulation. Model selection also showed that maternal environmental effects did not significantly explain variability in *Symbiodinium* communities found among larvae. This result, combined with the moderate heritability estimate, indicates that similarities in *Symbiodinium* communities among larvae of the same maternal brood were due to gene(s) inherited by these larvae.

Heritability estimates reveal important information about the evolvability of a trait, such as the capacity of brooding corals to vary their symbiont communities in response to changing environmental conditions. If levels of heritability and genetic variance are low, then responses to natural or artificial selection (evolvability) would be limited [48]. Conversely, high heritability and high genetic variance of a trait would enable greater responses to selection pressures. On the other hand, highly heritable symbiont communities with low genotypic variation could be problematic for vertically-transmitting coral populations if adult communities are thermally sensitive [50]. We found moderate heritability of *Symbiodinium* communities in *S. hystrix* (ShA). Much greater heritability of the *Symbiodinium* community was expected in this vertically-transmitting coral, especially in comparison with what is known of other important reproductive and fitness traits. For example, fertilization success, larval heat tolerance, protein content, settlement success, settlement substrate preferences, and juvenile growth and survivorship are all heritable traits [51–55]. Although the distribution of posteriors was skewed towards values greater than that of our heritability estimate, it is unlikely that heritability (i.e., genetic regulation) for this trait will resolve to be much greater with increased sampling effort (~0.5-0.6, Figure S2). The moderate levels of genetic regulation (i.e., heritability) found here suggest that *S. hystrix* (ShA) has some capacity to respond to changing environmental conditions. Thus, intervention efforts to facilitate such phenotypic change may be possible [48]. Given that assisted evolution efforts involving heat-selected *Symbiodinium* types show promise in horizontally-transmitting corals [56, but see 57], it may be that vertically-transmitting, brooding species with moderate fidelity like *S. hystrix* (ShA) could also be candidates for assisted *Symbiodinium* uptake.

### Combined maternal and environmental uptake produces locally adapted but flexible Symbiodinium communities

Detection of 93 larval-specific OTUs in this study demonstrates that brooding corals like *S. hystrix* (ShA) have a mixed-mode transmission strategy, in which dominant symbionts are transmitted vertically but additional background strains are acquired from environmental sources. Although adult diversity may have been under-sampled by only sequencing one branch of each parental colony, unique larval OTUs were not detected in any of the 45 adult colonies that were genotyped. Environmental uptake of novel *Symbiodinium* by larvae of this species is further supported by the appreciable amount of variation in the composition of larval *Symbiodinium* communities that was not under genetic control, according to our heritability model. These results validate the hypothesis of potential mixed-mode transmission initially raised by Byler et al. [20]. However, although juveniles hosted multiple symbionts in their study, they did not find differences in diversity between *S. pistillata* adults and larvae [20]. Evidence of mixed-mode transmission in *S. hystrix* (ShA) contradicts previous assumptions that maternally-transmitted symbiont communities are transferred to offspring with high fidelity in corals [23–25]. Our findings are consistent with transmission patterns documented in other symbiotic systems, such as wild *Drosophilia hydei* populations [9], *Acromyrmex* ants [15,16], and paramecium [17,18], and aligns symbiotic transmission ecology in corals with terrestrial invertebrate symbioses. Additionally, the novel diversity found in *S. hystrix* (ShA) larvae mirrors increased diversity of *Symbiodinium* communities detected in eggs of *Montipora capitata* and *M. digitata* compared to adults [21,22] and of bacterial communities in larvae of the brooding coral *Porites astreoides,* as well as of various bacterial communities associated with larvae of sponge species with supposed vertical transmission [31,58].

Mounting evidence for mixed-mode transmission across phyla suggests that it may be evolutionarily advantageous to compromise between completely vertically- and horizontally-acquired symbiont communities, as both strategies provide distinct advantages and disadvantages [14,20]. In *S. hystrix* (ShA), vertical transmission of *Symbiodinium* that are locally adapted to the parental environment is likely to provide benefits for a species that is able to self-fertilize [49,59] and has highly localised larval dispersal (e.g., 60–62]. However, a locally adapted community might become a liability if environmental conditions change or if larval dispersal distances are long. Negative effects include deregulation or disruption of symbiont abundances, which may have harmful physiological effects on the host, like bleaching in corals [38] or wasp parasitism in insects [9,63]. Thus, a mixed-mode strategy that results in superinfections of multiple symbionts can be beneficial [e.g., parasitoid protection in aphid hosts, 9,19] and may provide more flexibility for adjusting to variable environmental conditions. Similarly, a mixed mating strategy of selfing and outcrossing in *S. hystrix* (ShA), combined with a functional nutritional symbiosis upon release, may facilitate both local and long-distance dispersal [49]. Our findings confirm that diversity and flexibility in *Symbiodinium* transmission are greater than previously thought, highlighting the potential for evolvability that may confer greater resilience than coral species with strict vertical transmission.

Additional to environmental uptake of *Symbiodinium* during early ontogeny, processes like competitive exclusion may contribute to differences between larval and adult communities in *S. hystrix* (ShA). Theory suggests that competition among symbionts may preclude transmission of an exact replica of the parental symbiont community because conditions promoting growth for some symbionts may differ between life stages [11]. The novel symbiont diversity found in *S. hystrix* (ShA) larvae may provide benefits similar to those observed in insect symbioses, for example to provide larvae with the flexibility to host optimal symbiont types for the changing conditions through ontogeny [20,27]. For example, *Symbiodinium* C1_OTU136, which was uniquely identified in larvae, may represent an adaptive advantage for this early life stage. Clade C types are taxonomically and physiologically diverse [10,64], and exhibit a range of tolerances for light and temperature, which are also reflected in their *in hospite* distributions across individual adult colonies and species [65]. Larval settlement and early juvenile survival are generally highest in cryptic, low-light areas that offer protection from predation [66,67]. Given that optimal settlement environments differ substantially from light environments experienced by adults, potentially by as much as 10-fold [67], it is possible that variation in *Symbiodinium* communities between larvae and adults observed here relates to different selective pressures associated with differing light environments [68–70]. Other potentially numerous, uncharacterized differences between larval and adult microhabitats may also contribute to differences in selective pressures between life stages. The potential ecological roles for the larval-specific OTUs recorded here are unknown. Indeed, it is possible that they represent non-symbiotic, free-living types [71] that may have attached to the exterior of the larvae following release or that may have entered brooded larvae without engaging in symbiosis. Further work is needed to determine how many of these OTUs represent physiologically important versus transient *Symbiodinium.*

### *Potential mechanisms shaping larval* Symbiodinium *communities*

The immune system is an obvious mechanism by which the host could exert control over the symbiotic community by regulating the establishment of individual *Symbiodinium* types [72] or of either whole clades or functional units (i.e., clades or types with similar metabolic roles) [44]. The symbioses of *Wolbachia* and *Spiroplasma* bacteria among *Drosophila* and lepidopteran genera, for example, are highly specific and exclude other bacterial lineages through a dynamic and mature immune response, to the extent that specific *Drosophila* species host novel and specific *Wolbachia* and/or *Spiroplasma* strains [12,39]. Mechanisms of immunity that could be transmitted through inheritance of parental genes include components of both the innate and adaptive immune response, including some that have been implicated in shaping invertebrate symbiont communities, such as T-cells, Nod2, defensins, and antimicrobial peptides [as reviewed in 73,74]. These mechanisms have been documented during *Symbiodinium* establishment in corals [72,75,76] and observed in the Hydra/bacteria symbiosis [73,77].

Conversely, the greater variation and diversity found in larval compared to adult *Symbiodinium* communities may be a function of an immature immune response that is not yet able to differentiate appropriate *Symbiodinium* types, rather than an adaptive response. As the coral immune system matures over time [78,79], it is possible that a winnowing process eliminates symbionts that are not physiologically beneficial to the coral host [20,27]. If true, then the ubiquitous presence of *Symbiodinium* C1_OTU136 in larvae may be a consequence of an opportunistic *Symbiodinium* type taking advantage of immature host immunity. Further work is needed to identify the role that the immune response has in shaping *Symbiodinium* communities; in particular what (if any) immune-related genes are being transmitted from parents to offspring and whether novel symbionts are a function of an under-developed immune response.

### Winnowing and microhabitat variation may shape adult Symbiodinium communities

The disparate *Symbiodinium* communities in larvae versus adults found here further indicate that the re-shaping of the *Symbiodinium* community through ontogeny is an important developmental process in corals. Ontogenetic variability in microbial communities (both *Symbiodinium* and bacteria) is common in both vertically- and horizontally-transmitting cnidarian species [20,21,26,27,29–31,80–84]. The low level of variation in *Symbiodinium* communities associated with corals ranging in diameter from 8 cm to >30 cm [3-10 years; 85] suggests that the end of the winnowing process likely occurs earlier in the development of the brooding coral *S. hystrix* (ShA)(i.e., before 3 years) than in broadcast-spawning corals [~3.5 years; 26,27]. Although evidence for switching of symbiont communities in adults corals exists [86], the pre-winnowing period may be the most flexible time for hosts to associate with a diversity of microbes. Therefore, identifying at what stage winnowing occurs in brooding corals will provide crucial insights into when the flexibility to associate with environmentally-acquired and potentially stress-tolerant types diminishes and specialisation of the *Symbiodinium* community begins.

Spatial patterns in the abundances of *Symbiodinium* C120, D1, and D1a in adult corals were not consistent along or down the reef slope at Lizard Island, but may reflect variable temperature and light regimes at the microhabitat level that interact with differing photo-physiologies among symbiont types [87,88]. Variation in benthic light regimes at cm-level scales down the vertical faces of individual colonies and variation in irradiance within coral tissues have been shown to drive symbiont communities in other coral species [69,70,89,90]. Thus differences at the meter-level scales found in this study could be important for structuring *in hospite Symbiodinium* diversity, and may be partly responsible for the variability found at the level of individual larvae and broods. These small scale differences in symbiont abundances within adults may subsequently influence variability in *Symbiodinium* types among larvae. However, further work is needed to understand how fine-scale environmental variables impact *Symbiodinium* presence and abundance at the type and OTU level in this species.

## Conclusion

Based on novel heritability and paternity analyses, we show that *Symbiodinium* communities associated with the brooding coral *S. hystrix* (ShA) are only partially genetically regulated by their host and that larvae retain the flexibility to associate with novel symbionts across generations. Unexpectedly, our results reveal a mixed-mode transmission strategy for establishing *Symbiodinium* communities in larvae of a brooding coral, based on demonstrations that novel and unique *Symbiodinium* types are found in brooded larvae but not in adults. Importantly, this information aligns symbiosis transmission ecology in corals with well-known terrestrial invertebrate symbioses that typically exhibit mixed-mode transmission strategies. Advances in the understanding of heritable genetic mechanisms quantified here provide important insights into how *Symbiodinium* communities may be targeted for intervention strategies to increase reef resilience.

## Materials and Methods

### Study species and sampling design

The common, hermaphroditic coral *Seriatopora hystrix* broods sexually-produced larvae following internal fertilization of eggs by sperm from surrounding colonies [49,91]. DNA extracts of planula larvae for the present study were selected from samples that were collected in an earlier study to assess sperm dispersal distances and larval parentage of a cryptic species within the *S. hystrix* species complex, specified as *S. hystrix* (ShA) [49,92]. In Warner et al.’s study, colonies were tagged and sampled for molecular analyses within a 16 m x 16 m sampling area, with additional colonies sampled from two adjacent transects (totalling 16 m x 40 m area) in the Lizard Island lagoon [S14°41.248, E145°26.606; 49,92]. Microsatellite genotypes and paternity assigned to individual larvae in this earlier study [49] enabled us to examine the effect of both maternal and paternal identity on larval *Symbiodinium* communities across a full pedigree of larval relatedness. Hence, our study included full-sib and half-sib larvae, and four individuals produced by selfing (further details in Supporting information and Table S1).

### Symbiodinium community genotyping

*Symbiodinium* communities of adults and larvae were quantified with amplicon sequencing of the ITS-2 locus using the same DNA extractions that had been used to assign microsatellite genotypes and paternity in Warner et al. (2016). Nine maternal and 45 assigned paternal colonies (which included the nine maternal colonies), plus all larvae whose paternity was designated with a confidence level of Very High, High, or Medium by Warner et al. (2016) (n=60 larvae) were sequenced with the primers ITS2alg-F and ITS2alg-R [93] using paired-end Illumina Miseq technology. Library preparation and sequencing were performed at the University of Texas at Austin’s Genomics Sequencing and Analysis Facility (USA) using their standard protocols, including Bioanalyzer (Agilent)-based DNA standardization and pooled triplicate PCR before library preparation.

Raw reads (total = 6,875,177) were analysed using the USEARCH and UPARSE pipelines [v.7; 94], as outlined in Quigley et al. [95; further details in Supporting Information]. Because there is currently no single copy marker for *Symbiodinium* genotyping [96], the ITS-2 marker was selected for the broadest comparisons to the vast literature that has used this marker to describe *Symbiodinium* diversity, including some using next generation sequencing (e.g., [22,95,97–100]. Additional steps were taken to assess the presence and impact of intragenomic variants (further explained below). Briefly, reads were filtered, clustered into OTUs at 97% similarity, annotated with NCBI nt database and *Symbiodinium*-specific searches (further details in Supporting Information Table S2). Using these methods, the majority of the OTUs were re-assigned to a clade/type level, leaving only 0.03% of cleaned reads (1459 reads, 78 OTUs) that could not be classified, and which may represent new *Symbiodinium* types (Table S3, Figure S1, Supporting Information).

To account for variable read-depth across all samples, sample reads were normalized using ‘DESeq2’ and ‘Phyloseq’ implemented in R [101–103]. Nonmetric multidimensional scaling (NMDS) was performed and plotted using the normalized counts matrix using ‘Phyloseq’, ‘vegan’, and ‘ggplot’ [104,105]. Genetic distances between OTUs were calculated in ‘Ape’ [106]. Statistical testing of variation in OTU abundance was performed on raw reads in ‘DESeq2’, which incorporates variance normalization of OTU abundance, and interpreted using the Bejamini-Hochberg correction for multiple-inferences of p-adjusted alpha at 0.05. ‘DESeq2’ outputs are expressed in multiplicative (log2 fold) terms between or among treatments [107]. Network analysis on planula larvae was performed using the ‘igraph’ package [108] and custom scripts from [109].

### Estimating the diversity and heritability of Symbiodinium communities

To describe the *Symbiodinium* community in coral samples, we used a diversity measure (^q^D^Z^ _ij_(p)) that incorporates OTU richness, evenness and sequence similarity [110]. Sequence similarity was calculated using pairwise percent similarities between OTU sequences using the ‘Ape’ package with a “raw” model of molecular evolution. Heritability of *Symbiodinium* diversity associated with the 60 larvae was calculated using the package ‘MCMCglmm’ [111] utilizing the diversity metrics described above, where the coefficient of relatedness between individuals was set as a random effect. Models were run with 1.5 × 10^6^ iterations, a thinning of 50, and burn-in of 10% of the total iterations. A non-informative flat prior specification was used following an inverse gamma distribution [112]. Assumptions of chain mixing, normality of posterior distributions, and autocorrelation were met. The posterior heritability was calculated by dividing the model variance attributed to relatedness by the sum of additive and residual variance. Deviance Information Criterion was used to test if adding a maternal random effect had a statistically significant effect on heritability estimates.

### Multiple ITS-2 copies and intragenomic variation

Intragenomic variation within and between *Symbiodinium* types makes classifying type-level diversity in *Symbiodinium* based on sequence data difficult [99,113–115]. However, comparisons between single-cell and next-generation sequencing suggests that clustering across samples at 97% similarity sufficiently collapses intragenomic variants to the type level [99], as has been used in this study. Furthermore, a recent study suggests that clustering across samples at 97% identity underestimates diversity instead of overestimating it [109]. Intragenomic variation and generation of false-positives is therefore substantially minimized by using across-sample clustering at 97% similarity, as we have employed. Single cell sequencing is currently financially and logistically outside the scope of studies that examine communities of hundreds of different *Symbiodinium* types (as with coral juveniles), with a majority of these types not yet existing in culture. It is questionable if microsatellite flanking regions provide superior taxonomic resolution [116], and as no known single-copy marker exists, using other markers in tandem with ITS-2 will only result in data representing multiple, multi-copy markers. We addressed intragenomic variation by clustering across samples at 97% similarity and also provided two additional analyses to test for their presence and potential impact on the heritability estimate; and both confirm the robust nature of our conclusions in regards to this issue. Overall, we undertook a three-step approach, as outlined in [95], to assess if multiple copies and intragenomic variation of ITS-2 genes could potentially bias abundance and heritability estimates across *Symbiodinium* types after clustering at 97% identity. Briefly, OTUs were first divided by clade and inspected for co-occurrence across samples using the tree function in ‘Phyloseq’ and grouped into subsets of co-occurring OTUs. Secondly, OTUs that increased proportionally and with high percent pairwise similarity were inspected. Finally, pairwise percent identities were calculated for these latter subsets of OTUs using the package ‘Ape’ [106], and correlations of variance-normalized abundances were calculated for pairs that had greater than 85% similarity with the function ggpairs in the package ‘GGally’ [104]. The diversity metric was calculated taking into account possible intragenomic variation by pooling the raw abundances of potential intragenomic variants (OTUs: 8/10, 12/22/24, 28/223, 3/6, 588/848), and heritability was calculated using the parameters described above. As we found little evidence of intragenomic variation amongst OTUs, these results and their impact on heritability estimates are only discussed in the Supporting Information.

### Colony size and spatial distribution of adult S. hystrix (*ShA*) colonies

To determine if *Symbiodinium* communities varied with colony size (as a proxy for colony age), adult colonies were divided into five size classes based on their mean diameter [49]: < 8 cm (n = 1 colony), 8 – < 14 cm (n = 19), 14 – < 20 cm (n = 13), 20 – < 26 cm (n = 11), and 26 – 32 cm (n = 1). Differential abundance testing of *Symbiodinium* OTUs was among size classes was performed as for larval communities.

Sitepainter [117] and Inkscape [118] were used to test for spatial patterns in the distribution of *Symbiodinium* OTUs associated with the 45 adult colonies of *S. hystrix* (ShA) that were genotyped across the 16 m x 40 m sampling area. Gradient Boosted Models and linear models were run in the package ‘gbm’ [119] to examine spatial distributions of the ten most abundant OTUs. Linear models were checked for assumptions of linearity, normality, and homogeneity of variance. Square-root transformations were used to correct for issues of normality or heterogeneity. Latitude and longitude coordinates were centered before fitting models. The package ‘Spatstat’ [120] was used to visualize spatial variability in abundances of the three most significantly heterogeneous OTUs across the sampling area (OTUs: 1, 3, and 6). Spearman’s Rho rank correlation coefficients were calculated to test for competitive exclusion amongst the three OTUs that varied significantly across the sampling area. Pairwise p-values were generated for all OTU comparisons using the base ‘stats’ package in R.

## Acknowledgments

This study was carried out with permission and in accordance with the recommendations from the Great Barrier Reef Marine Park Authority.

## Data Accessibility

DNA sequences: All sequencing data will be made available through NCBI SRA.

## Author contributions

K.M.Q. and B.L.W. conceived of the experiment. K.M.Q. and P.A.W. designed the sampling scheme and performed the experiment, collected and analysed the data. L.K.B. provided reagents and materials. K.M.Q. wrote and B.L.W., L.K.B., and P.A.W. edited and critically reviewed the manuscript. All authors read and approved the final version.

